# The connectome spectrum as a canonical basis for a sparse representation of fast brain activity

**DOI:** 10.1101/2021.03.03.433561

**Authors:** Joan Rué-Queralt, Katharina Glomb, David Pascucci, Sebastien Tourbier, Margherita Carboni, Serge Vulliémoz, Gijs Plomp, Patric Hagmann

## Abstract

The functional organization of neural processes is constrained by the brain’ s intrinsic structural connectivity. Here, we explore the potential of exploiting this structure in order to improve the signal representation properties of brain activity and its dynamics. Using a multi-modal imaging dataset (electroencephalography, structural MRI and diffusion MRI), we represent electrical brain activity at the cortical surface as a time-varying composition of harmonic modes of structural connectivity. The harmonic modes are termed connectome harmonics, and their representation is known as the connectome spectrum of the signal. We found that: first, the brain activity signal is more compactly represented by the connectome spectrum than by the traditional area-based representation; second, the connectome spectrum characterizes fast brain dynamics in terms of signal broadcasting profile, revealing different temporal regimes of integration and segregation that are consistent across participants. And last, the connectome spectrum characterises fast brain dynamics with fewer degrees of freedom than area-based signal representations. Specifically, we show that with the connectome spectrum representation, fewer dimensions are needed to capture the differences between low-level and high-level visual processing, and the topological properties of the signal. In summary, this work provides statistical, functional and topological evidence supporting that by accounting for the brain’ s structural connectivity fosters a more comprehensive understanding of large-scale dynamic neural functioning.

## Introduction

The brain is a large biological network of interconnected neural populations. Given the vast number of neural populations, at the macroscopic level, the activity of the brain is commonly studied using a parcellation. The choice of brain parcellation into areas of interest reflects the underlying assumption on how large populations of neurons cluster together, and typical clustering models are anatomical (*1*), functional (*2, 3*), structural (*4*) or multi-modal (*5*). Considering the entire brain, brain activity can be represented as a trajectory over time in a high-dimensional coordinate system where each dimension represents the activity of a specific brain area. Even though these brain areas are commonly regarded as independent units, they synchronize among them according to their connectivity (*6*). In this work, we focus on local measures of cortical electrical activity as measured by source reconstructed electroencephalography (EEG). We refer to the spatial organization of the brain activity signal as its underlying structure, which takes the form of a graph (see Fig. S1 for illustration). The underlying structure of the signal allows us to draw observations from a global perspective (as opposed to local). For example, in a movie, the underlying structure is defined by a two-dimensional grid, in which pixels are connected to their four nearest neighbours. This underlying structure allows us to determine the relevance of a pixel value (signal) at a given time-point, i.e., data from neighbouring pixels can be seen as redundant, if both pixels encode a piece of background, or very significant, if they are forming an edge belonging to a contour. In the case of the brain, the underlying structure is defined by the structural connectome (SC) (*7, 8*). The SC can be conceptualized as a graph in which the nodes are defined as gray matter brain areas using a reference atlas (see Fig. 1C), and the edges reflect some properties of the estimated white matter tracts connecting each pair of brain areas (number of fibers, their average length or mean fractional anisotropy), which are estimated using diffusion magnetic resonance imaging (*9*) (dMRI, see Fig. 1D.).

**Fig. 1.**
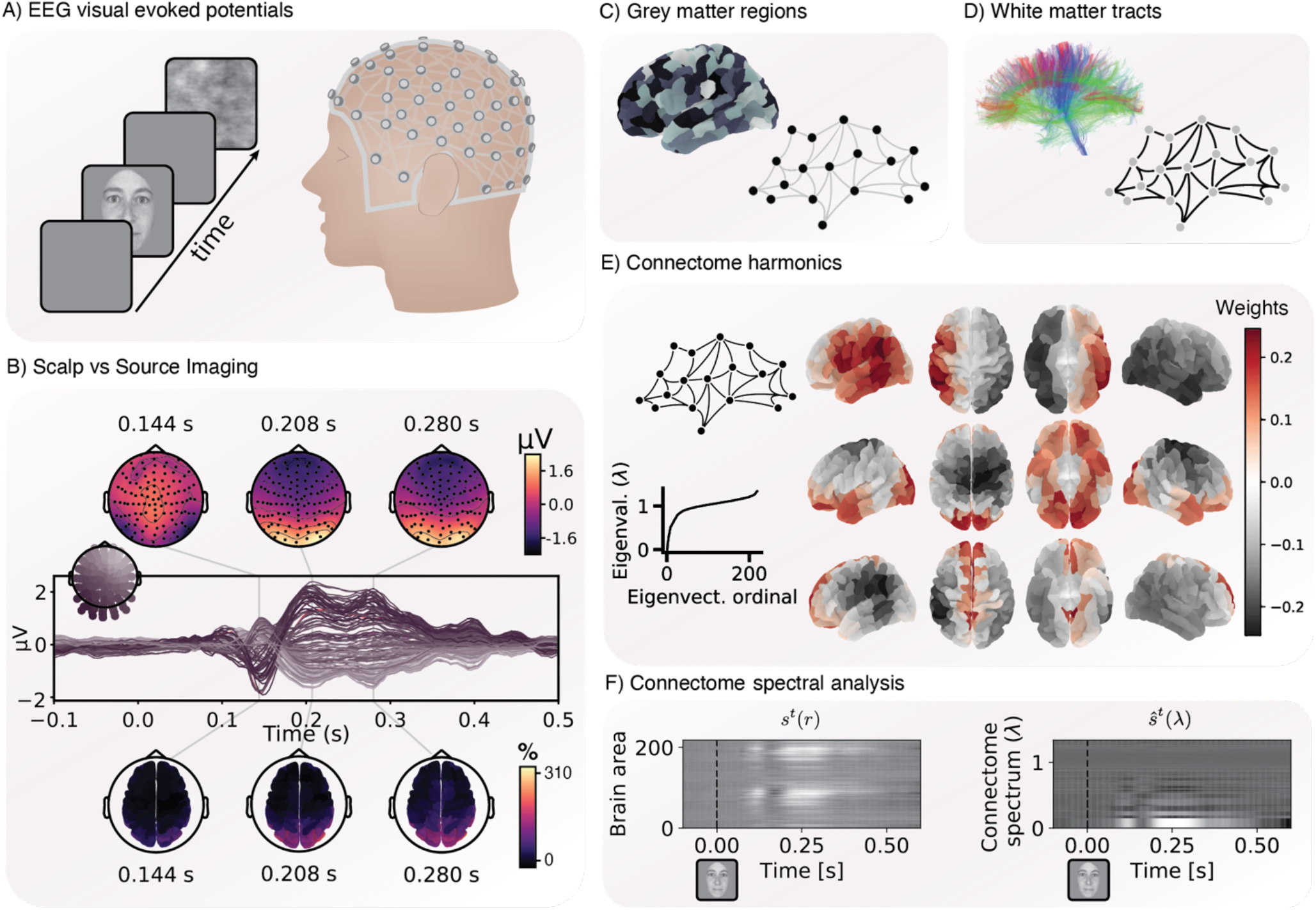
Connectome harmonics for visual evoked potentials. **(A)** The visual evoked potentials’ experimental design consisted in the presentation of two types of stimuli: faces and scrambled-faces. The face images were taken from an openly available dataset (*52*) and cropped with a Gaussian kernel to smooth the borders. For each one of the 19 participants, approximately 200 trials for each condition were recorded and aligned at the onset of the stimulus presentation. **(B)** Scalp patterns before, and after the face stimulus is presented (at 0 ms) (generated with the visualization tool of MNE v0.21), and the source reconstruction of the stimulus evoked activity. **(C)** Grey matter areas defined by an anatomical parcellation define the nodes of the brain graph. **(D)** The estimated white matter tracts from diffusion MRI define the connectivity strength of the edges of the graph. **(E)** From the constructed graph, we perform the graph Laplacian eigendecomposition and obtain a graph spectrum, consisting on a set of eigenvalues and a set of eigenvectors, the former ones termed graph spectrum, and the later termed graph or connectome harmonics. A few more examples of connectome harmonics are shown in Fig. S1. **(F)** The source reconstructed time-series signal represented in each brain area r (left plot, S^E^(r)) or in each connectome spectrum corresponding to each graph Laplacian eigenvalue λ (right plot, ŝ^*t*^(λ)).

When analyzing a signal of which the underlying domain structure is known and can be defined by a graph, such as the structural connectivity, we can decompose the signal as the sum of graph Laplacian eigenvectors by means of the graph Fourier transform (*10*) (GFT). In the case of the brain, we define connectome spectral analysis as the connectome graph Fourier Transform of the brain activity signal:

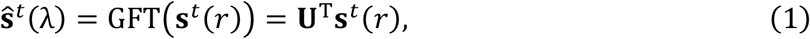

where **s**^*t*^(*r*) represents the activity of a specific brain region *r* at time *t*, and the columns of *U* contain the connectome Laplacian eigenvectors {**u**_i_}, from now on referred as connectome harmonics (*11*). The signal’ s connectome spectrum ŝ^*t*^ (λ) defines the participation of each connectome harmonic in the signal **s**^*t*^ (*r*) (*12, 13*) (see Fig. 1E-F and Fig. S1). GFT is analogous to the two-dimensional Fourier transform for images, for which the domain is a regular grid, or to the discrete one-dimensional Fourier transform for temporal signals, for which the domain is a discrete circle. Equivalent to how sinusoidal Fourier modes (Laplacian eigenvectors) relate to frequencies (Laplacian eigenvalues) in the one-dimensional Fourier transform, connectome harmonics {**u**_*i*_} are also ordered by their associated connectome graph Laplacian eigenvalue λ_*i*_ which quantifies their smoothness (in terms of Dirichlet energy (*14*)). We will refer to the set {λ_*i*_} as connectome spectrum in the rest of the manuscript. Low-frequency connectome harmonics capture smooth signal gradients over the connectivity graph, for example capturing the left-right, anterior-posterior, ventral-to-dorsal and medial-peripheral axes as interconnected (*11*). By contrast, high-frequency harmonics capture irregular patterns over the connectivity graph in space (see Fig. S1 for illustration).

The structure-function relationship is an unresolved topic in neuroscience (*6, 15, 16*), and recently, several studies have explored potential applications of connectome spectral analysis (see (*17*) for a review). Characterizing brain function by means of its underlying structure is motivated by the fact that brain activity is constrained by, and coupled through its underlying connectivity structure (*18*). When functional interactions follow structural connections, as in the case of brain activity, the signal is represented in the smoothest subset of connectome harmonics, and thus, it can be reconstructed with a sparse signal representation (*13, 19*). Using the underlying structure of the signal might not only improve the statistical properties of the signal, but it could also provide mechanistic information on how signals propagate. In this sense, connectome harmonics have been theoretically proposed as a mechanism for macroscopic brain activity, allowing nested functional segregation and integration across multiple spatio-temporal scales (*20*).

Even though connectome harmonics seem a promising tool for studying whole brain activity, a comprehensive empirical evaluation of the properties of the signal’ s connectome spectrum is lacking. In this work, we characterized the advantages of connectome harmonics as a coordinate system for representing the fast-evolving brain-wide activity signals at the cortical surface estimated from EEG recordings. We report some evidences suggesting the advantages of the connectome spectrum representation in three different aspects: the statistical, the functional, and the topological properties of the signal. We first show that during visual evoked brain activity, the EEG signal estimated at the cortical surface can be represented more compactly by its connectome spectrum than with traditional atlas-based signal representation. This model-based representation of the signal performs equally or better in terms of compactness than traditional data-driven approaches such as PCA and ICA. Importantly, the compactness of the signal’ s connectome spectrum is specific to the connectivity structure, rather than to the graph spectral decomposition properties. Then, we demonstrate that the representation of the signal on its connectome spectrum automatically characterizes brain activity dynamics in terms of the signal broadcasting status, revealing integration and segregation regimes of brain processing, which follow consistent dynamics across participants. Finally, we provide evidence indicating that connectome harmonics capture functional aspects of visual perception with fewer degrees of freedom than isolated brain areas. These advantages extend to topological properties of the signal and their fast temporal dynamics. After presenting these three types of evidence, we propose the connectome spectrum as a canonical basis for the representation of large-scale brain activity dynamics.

## Results

### A sparse basis for large-scale brain activity

Coordinate systems in which data are represented compactly are advantageous because they summarize the data well, and they provide robustness to small variations such as noise (*21*). Furthermore, if brain activity admits a sparse signal representation, it means that the process has a limited number of parameters that we are able to measure. The compactness of the signal indicates how compressible it is, and gives us an estimation on its sparsity (*22*). The brain activity signal was recorded in visual evoked potential experiments using high-density EEG, and reconstructed at the cortical surface (see Fig. 1A,B). Using an anatomical parcellation, the single-trial cortical time-courses were parcellated into 219 areas (*23*) using the method introduced in (*24*). We estimated the sparsity of the brain activity in the usual brain atlas-based signal representation, in which the signal is represented as a set of data-points in a coordinate system spanned by the set of brain areas *r*_*i*_, (i.e., **s**^*t*^(*r*)). Using the structural connectivity, we computed the GFT (*10*) of the signal **s**^*t*^(*r*) to obtain its connectome spectrum ŝ^*t*^(λ) (*13*). Then we compared the compactness of the signal **s**^*t*^(*r*) to the compactness of ŝ^*t*^(λ).

The compactness of the evoked signal in each coordinate system was quantified by means of signal compression performance (inspired from (*25*)). Signal compression was performed by replacing the signal values (and its spectrum coefficients) with magnitudes smaller than the *p*-th percentile with zeros (and performing the corresponding inverse GFT on the resulting coefficients in the case of the signal connectome spectrum). The compactness for each *p* was then measured as the Pearson correlation between the original and the compressed signal on the one hand, and as one minus the normalized mean squared error (NMSE, see section *Methods: Analysis of Compactness*, Fig. 2A,B) on the other hand. Results show that thresholding performed in the connectome spectral domain achieved greater signal compactness than in the atlas-based coordinates (*p* <.001, Wilcoxon Rank-sum two-sided test, corrected for multiple comparisons via Bonferroni, Fig. S2A,B). This indicates that ŝ^*t*^(λ) is sparser than **s**^*t*^(*r*), meaning that the evoked activity is described with fewer connectome harmonics than brain areas. The idea that the connectome spectrum of the signal is sparse is illustrated by the fact that we can reconstruct more than 50% of the explained variance of the signal after compressing down to just 5% of its connectome spectrum content. For validation, we tested if the improved compression capacity was related to the representation of the signal in its underlying structure, and not to general decomposition properties of the graph Laplacian. To this end, we performed a non-parametric statistical analysis by measuring the compactness of the signal represented by degree-preserving surrogate connectome harmonics (see *Methods: Surrogate harmonics*). The compactness of the signal’ s connectome spectrum was significantly higher than in the spectrum defined by degree-preserving surrogate graphs, in terms of signal correlation and MSE (*p*<.001, Wilcoxon Rank-sum two-sided test, corrected for multiple comparisons via Bonferroni, see Fig. 2B, Fig. S2A,B). In fact, both metrics of compactness performance in the surrogate harmonics were not significantly different to the ones of the **s**^*t*^(*r*) signal *p*>.05). This shows that the actual structure of the connectome drives the observed compression.

**Fig. 2.**
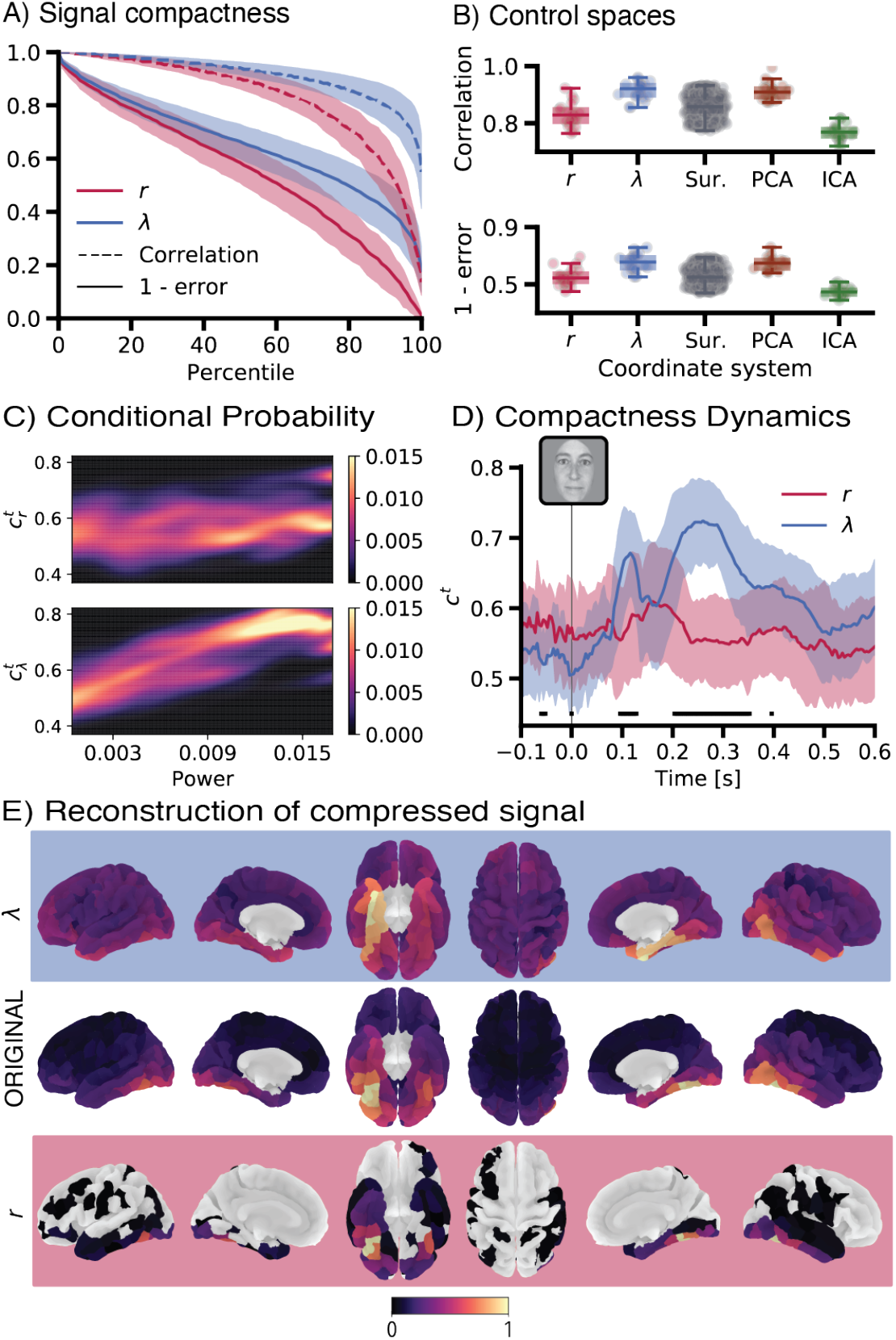
Signal compactness in different coordinate systems of signal representation. **(A)** Signal compactness measured by two distance metrics between original and compressed signals: correlation (dashed line) and reconstruction error (continuous line) (see *Methods: Analysis of compactness*). The signal compression was performed at all integer percentiles from 1 to 100 for the signal represented by brain areas (*r*) and connectome harmonics (*λ*). **(B)** Signal average compactness (mean across percentiles). It was computed for different controls (see *Methods*): Surr., surrogate connectome harmonics; PCA, principal components from principal component analysis; and ICA, independent components from independent component analysis. See Fig. S2 for statistical significance tests of the differences among coordinate systems. **(C)** Conditional probability of the compactness dynamics *c*^*t*^ (shown in **(D)**). It is conditioned by the power of the signal at the same time-point (see *Methods: Conditional probabilities*) for either the area and the connectome harmonics’ activity. **(D)** Compactness dynamics *c*^*t*^. It is computed at each time-point as the compression reconstruction performance (1 -reconstruction error) averaged across percentiles (see *Methods: Analysis of compactness*), and informs about the number of active dimensions in time (i.e. a proxy of its sparsity). At each time-point, a Wilcoxon Rank-sum two-sided test is performed, and those time-points with significant *p*-value are indicated with black rectangular boxes (Bonferroni corrected for multiple comparisons). **(E)** Visual illustration of the effect of compressing a signal. The signal is compressed at 95-th percentile in each coordinate system (only 11 out of 219 kept dimensions). For the atlas-based signal representation, the signal is shown after being compressed. For the connectome spectrum of the signal, the compressed activation coefficients are projected to the original atlas-based coordinates. The error bars show the mean and standard deviations of the distribution across subjects.

It is worth noting that the connectome spectral decomposition given by the connectome GFT is model-driven (we model brain activity as a weighted sum of structural connectivity gradients). Usually, signal decomposition techniques are data-driven (based on the signal properties), such as in the case of Principal Component Analysis (PCA) or Independent Component Analysis (ICA). Fig. 2B assesses the compression properties of these well-known methods to benchmark the performance of connectome harmonics. It shows that connectome spectral analysis compares similarly to PCA in terms of correlation and reconstruction error, but it always outperforms ICA (see Fig. S2C,D; *p*<.001, Wilcoxon Rank-sum two-sided test, corrected for multiple comparisons via Bonferroni). These results suggest that connectome harmonics capture well the variance of stimulus-evoked brain activity, and more specifically, that they compare favorably with commonly-used data-driven decomposition methods.

### Compactness dynamics

Visual evoked potentials have a highly temporal non-stationary nature; therefore, the sparsity of the signal can be expected to change in time. For this reason, we assessed the dynamical properties of the signal in the coordinate system defined by the connectome harmonics. We hypothesized that the fast dynamics of the visual evoked activity should be reproducible across participants if the signal’ s connectome spectrum (ŝ^*t*^(λ)) would capture relevant features. We suggested to use the compactness dynamics (*c*^*t*^), defined as the reconstruction error of the compressed signal at a given time-point, averaged across all percentiles (*Methods: Analysis of compactness*), as a proxy of the number of active dimensions at each time-point, i.e., the sparsity of the signal (Fig. 2D).

We first studied the dependency of the connectome spectral compactness on the power of the signal, to assess whether the activation of connectome harmonics was not trivial. We analyzed the relationship between the signal compactness and the total energy of the signal by means of the conditional probability *P*(*c*^*t*^ | ‖ · ‖_2_) (*Methods: Conditional probabilities*), for “·” representing **s**^*t*^(*r*) or ŝ^*t*^(λ). The conditional probability showed a strong linear tendency for the connectome spectrum ŝ^*t*^(λ) (see Fig. 2C), with an increase in the power of the signal positively correlated to the activation of a few connectome harmonics. This result suggest that connectome spectrum compactness is non-randomly distributed in time, and can be partially explained by the dynamics of the power of the signal. In other words, in moments of high power the signal is concentrated in few harmonics. When analyzing the original signal **s**^*t*^(*r*), however, we did not find such a relationship, indicating that signal power does not depend on the number of active areas.

To test whether connectome harmonics can characterize fast, functionally specific processes, we used visual-evoked potentials (VEP) to faces. Face-specific processes are well-studied and can be primarily localized to the Fusiform gyrus (*26*), but are known to also involve occipital, parietal and orbitofrontal areas (*27*–*29*). They also have a specific temporal signature in terms of temporal localization (at around 170 ms after the stimulus is presented) (*30, 31*).

The compactness dynamics of connectome harmonic 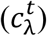 in the evoked brain signal revealed different regimes of visual processing (Fig. 2D). At time zero (0 ms), the visual stimulus was presented and the brain signal associated to stimulus processing quickly built up, becoming more compact in both the connectome harmonic and atlas-based signal representation. This increased compactness was significantly larger for the connectome spectrum of the signal than for the atlas-based signal in three time-windows ([96 – 128 ms, 204 – 352 ms, 396 – 400 ms], *p*<.05, Wilcoxon Rank-sum two-sided test, corrected for multiple comparisons via Bonferroni). During these time windows, the variance of the signal was captured by fewer number of harmonics than number of brain areas, suggesting that the constitution of the signal during those time-points is shaped by the activity of a limited number of structural networks rather than the activity of single brain regions. During pre-stimulus time, the signal reflects average activity across many non-aligned trials, and can be regarded as residual noise, evenly distributed across dimensions. For this reason, pre-stimulus time was not compressible in any coordinate system. Such noise was more evenly distributed in the signal’ s connectome spectrum, and thus, significantly less compact (*p*<.05, Monte-Carlo simulations, corrected for multiple comparisons via Bonferroni).

At this point, the reader might think that a compact signal might misses crucial information. To guarantee that the increased compactness of the signal does not come at the expense of destroying parts of the signal that are known to be relevant, we next focused only on two well-known spatio-temporal patterns: face processing in the ventral stream at 170 ms, and motion processing in the dorsal stream at 150 ms (*Methods: EEG*). For simplicity, reconstruction error and correlation after compression were estimated for three visual systems: ventral, dorsal and the group of early visual areas (see Fig. S3). The only significant differences in compactness properties between **s**^*t*^(*r*) and its connectome spectrum ŝ^*t*^(λ) were found in motion perception at 150 ms (*p*<.05, corrected for multiple comparisons via Bonferroni), in which compactness was better in the connectome spectrum ŝ^*t*^(λ). These results indicate that signal compression in the connectome harmonic representation does not remove the part of the signal known to be functionally relevant.

### Integration and segregation dynamics during visual perception

Low-order harmonics (i.e, the connectome Laplacian eigenvectors associated with the smallest eigenvalues) define smooth gradients of structural connectivity in brain space. They can be understood as patterns of integrated activity, i.e., brain activity in which sets of neighbouring brain regions show strong long-range coupling (in graph space). Conversely, high-order harmonics reflect short-range coupling, i.e., capture signal similarity between smaller sets of neighbouring areas, and in this sense reflect segregation mechanisms (*17*). During cognitive processes, brain activity shows a strong opposing relationship between the participation of these two types of cortical gradients (*32*), suggesting two different broadcasting mechanisms underpinning functional dynamics (*33*). In line with these ideas, we studied the dynamics of energy spread between the lower and higher end of the spectrum. Fig. 3A (and Fig. S4A for a different visual experiment) shows that the signal’ s power spectral distribution on the graph has a 1/λ shape. This means that most of the power is concentrated in spatial patterns that are smooth (dominance of low frequencies). To investigate the differential broadcasting dynamics of the low and high frequency parts, we split the graph spectrum into two parts, following the methodology introduced in (*34*) (see *Methods: Graph spectrum dichotomization*). We computed the ℓ_2_ norm of the part of the signal belonging, respectively, to the low-order connectome harmonics, and to the high-order connectome harmonics:

**Fig. 3.**
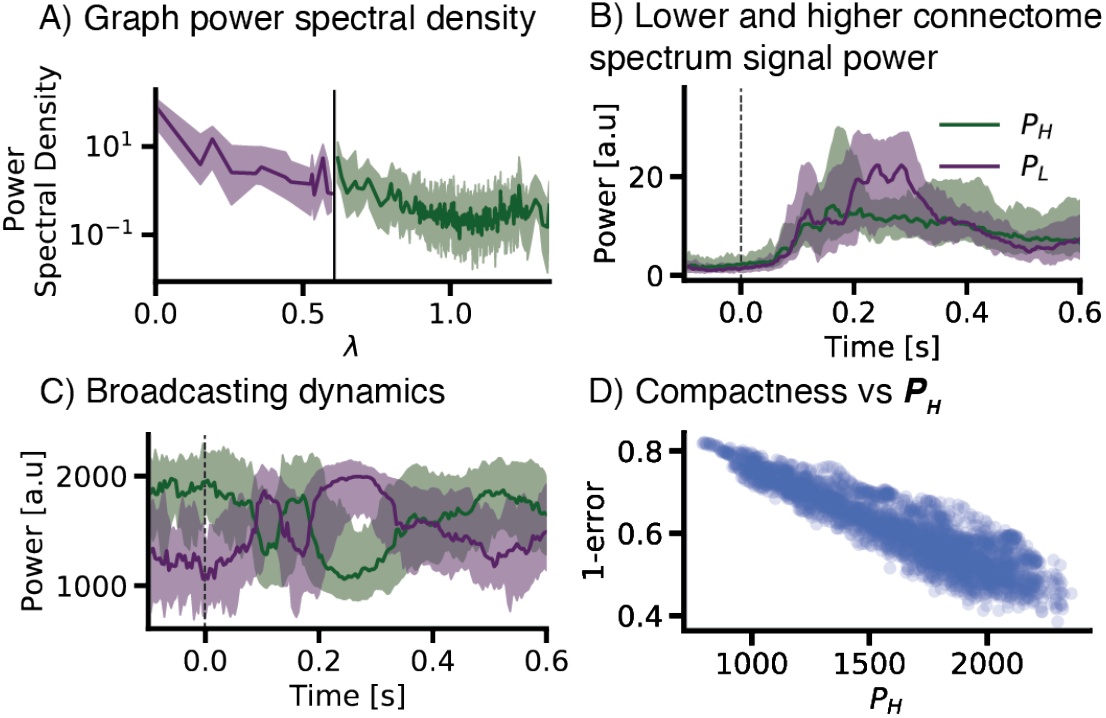
Signal broadcasting profile. **(A)** Graph power spectral density. It was computed as the power in each connectome harmonic during the whole time-series (see *Methods: Graph spectrum dichotomization*). The power spectral density was then divided into two halves of equal cumulative power. **(B)** Energy of the original signal contained in each of these two ends of the connectome spectrum. It was computed for each time-point: 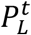, associated to integration processes, and 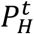, associated to brain activity segregation, respectively. **(C)** Time-courses in **(B)**. They were normalized by the total power of the signal, illustrating the normalized contribution across time to the total power of the signal. By definition, these time courses automatically profile the brain activity broadcasting profile into integration and segregation dynamics. D) The normalized 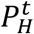 (segregation) at each time-point (shown in **(C)**) was compared to the connectome spectrum compactness at each time-point (here indicated as 1-error, previously shown in Fig. D). The continuous lines and the shaded area in the plots of this figure show the mean and standard deviations of the distribution across all subjects.

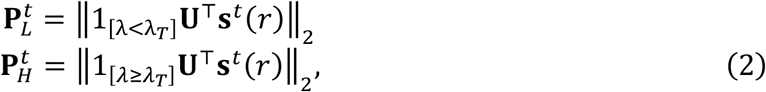

where 1_[λ<λ*T*_] is the indicator function, which keeps only the lowest part of the spectrum. The dynamic interplay between 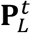 and 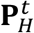 power change in time is shown in Fig. 3B,C, (also in Fig. S4B,C for a validation in a different dataset) and is strongly related to the compactness of the signal in the connectome harmonics coordinate system, as shown in Fig. 3D. There is a strong linear dependency of the dynamic evolution of the interplay between 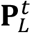 and 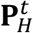 and the compactness of the connectome spectrum, indicating that during time-points at which compactness is maximal, brain activity is coupled over long-range network distance (integrated). In other terms, almost every time that the connectome spectrum is compact, the energy of the signal is concentrated in the low-frequency harmonics. We refer the reader to the *Discussion* section for a detailed interpretation on the obtained dynamics of integration and segregation.

### Low dimensional trajectories during visual perception

To understand how brain activity spans a given coordinate system, it is useful to conceive of the activity as a dynamic trajectory through that space (commonly referred as state-space representation). This kind of signal representation facilitates understanding whether some dimensions of the coordinate system are more important than others. Here, were compared activity trajectories evoked by images of faces to the trajectories evoked by images of scrambled-faces (see Fig. 4A), both when the coordinates represent the activity of brain areas (**s**^*t*^(*r*)), or the participation of connectome harmonics (ŝ^*t*^(λ)). Scrambled-faces are a commonly used control for the influence of low-level stimulus features, preserving an equal amount of image contrast and light intensity as in face images. The trajectories in Fig. 4A show that the activation of the different dimensions over time are not completely independent in any coordinate system: the mean Pearson correlation among dimensions in time was 0.54 for brain areas and 0.45 for connectome harmonics (*p*<.005, Wilcoxon Rank-sum two-sided test). However, ŝ^*t*^(λ) was significantly less co-linear than the signal **s**^*t*^(*r*) (see Fig. 4B, *p*<.005, Wilcoxon Rank-sum two-sided test). These results indicate that connectome harmonics capture components of brain activity that are more linearly independent in time, hence more likely to capture its degrees of freedom.

**Fig. 4.**
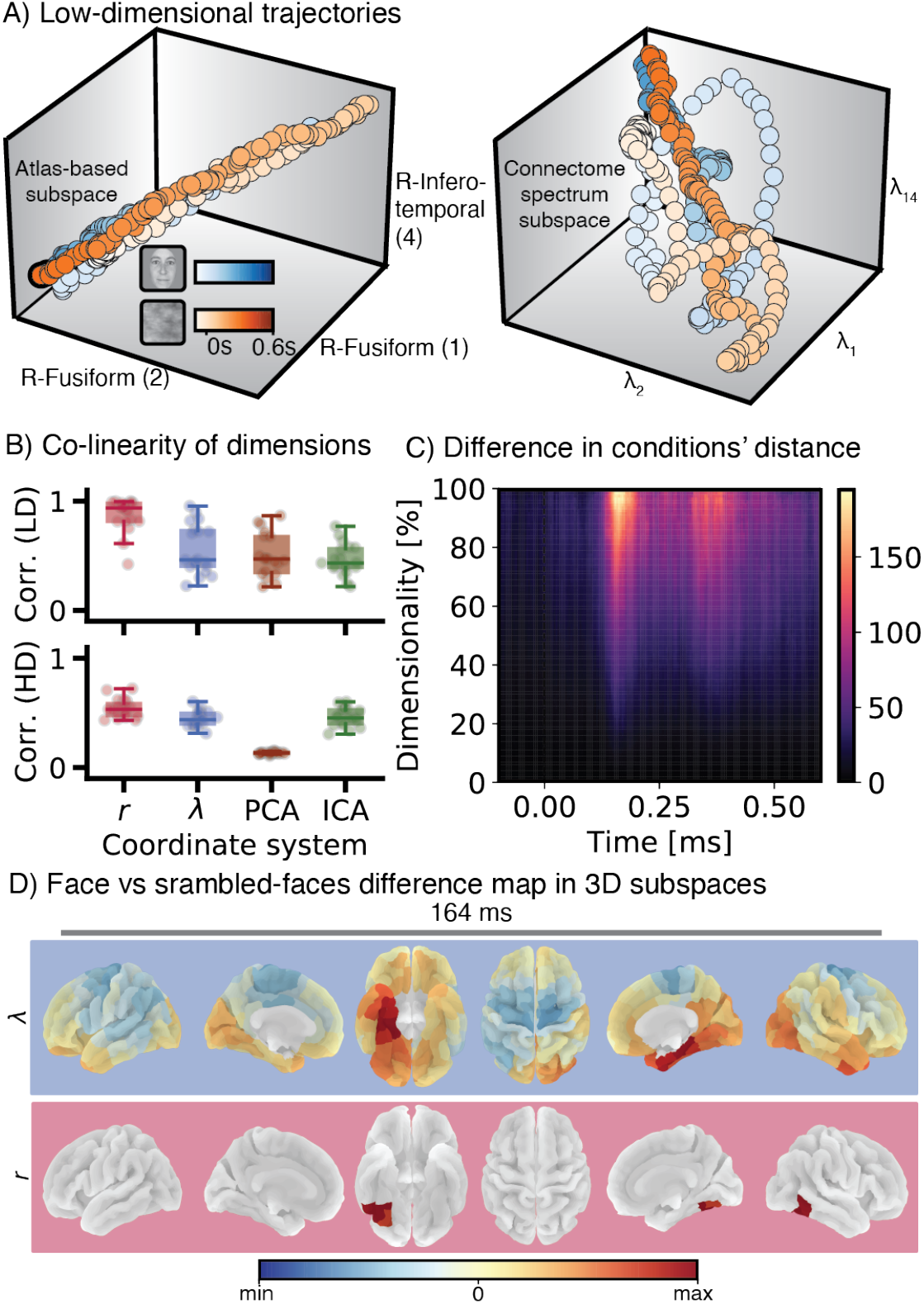
Trajectories of brain activity. **(A)** Low-dimensional trajectories. The brain activity signal evoked by stimulation with images of faces (in blue) and scrambled-faces (in red) is represented as a vector moving in time in a three-dimensional coordinate system. The three-dimensional space is defined by those dimensions that capture more energy of the signal: (left plot) coordinate system spanned by the three top brain areas, and (right plot) coordinate system spanned by the three top connectome harmonics. **(B)** Co-linearity of dimensions. The co-linearity of the dimensions of each coordinate system (*r*: atlas-based; λ: connectome harmonics; PCA: principal components from principal component analysis; ICA: independent components from independent component analysis) is measured by the average pairwise correlation among the top three dimensions (top plot), and among all dimensions (bottom plot). **(C)** Difference in conditions’ distance. The difference between the magnitude of the vector that connects the trajectories of the two conditions at each time-point is plotted for different percentile of compression (see *Methods: Distance between two trajectories*). **(D)** Face vs scrambled-faces difference map. The reconstructed pattern (for a compression percentile of 99, corresponding to keeping only the 3 dimensions shown in **(A)**) at the time-point in which the distance between the faces’ and the scrambled-faces’ trajectories is maximal in the atlas-based full dimensional signal (219 areas).

The degrees of freedom that encode functionally relevant features in a signal are expected to have an important proportion of the signal’ s energy. Here, we projected the signal into the few dimensions that concentrated most of the energy (by means of ℓ_2_-norm) to estimate the quality of different coordinate systems (see *Methods: Low-dimensional embedding*). First, we observed a larger distance between the two stimulus trajectories when projecting the signal in the connectome spectrum, compared to the signal defined by brain areas. This effect is illustrated in Fig. 4C, and demonstrates that, as the dimensionality of the coordinate systems gets reduced, the two trajectories become more separable in the signal connectome spectrum. Furthermore, the difference vector in the reduced subspace of the signal’ s connectome spectrum encodes a more representative brain activity map of the face-processing network (*27, 28*) (see Fig. 4D). In other words, the connectome harmonics that concentrate most of the stimulus-evoked energy capture the differences between faces and scrambled-faces with higher sensitivity than the area-based analysis. To put it differently, high-level features of visual stimulus are encoded by a smaller number of connectome harmonics than brain areas. This result supports the idea that connectome harmonics are able to capture independent neurophysiologic parameters.

Another important property of the state-space representation is that potentially functionally relevant features of the signal are encoded by the signal trajectory (*35*). Accordingly, we measured for each system of coordinates how the dimensions with highest energy contributed to the overall signal trajectory using tools from Topological Data Analysis (*36, 37*). Persistence homology of simplicial complexes allowed us to summarize the shape of the signal point cloud, and respectively its spectrum, by a persistence diagram (see Fig. 5A-C).

**Fig. 5.**
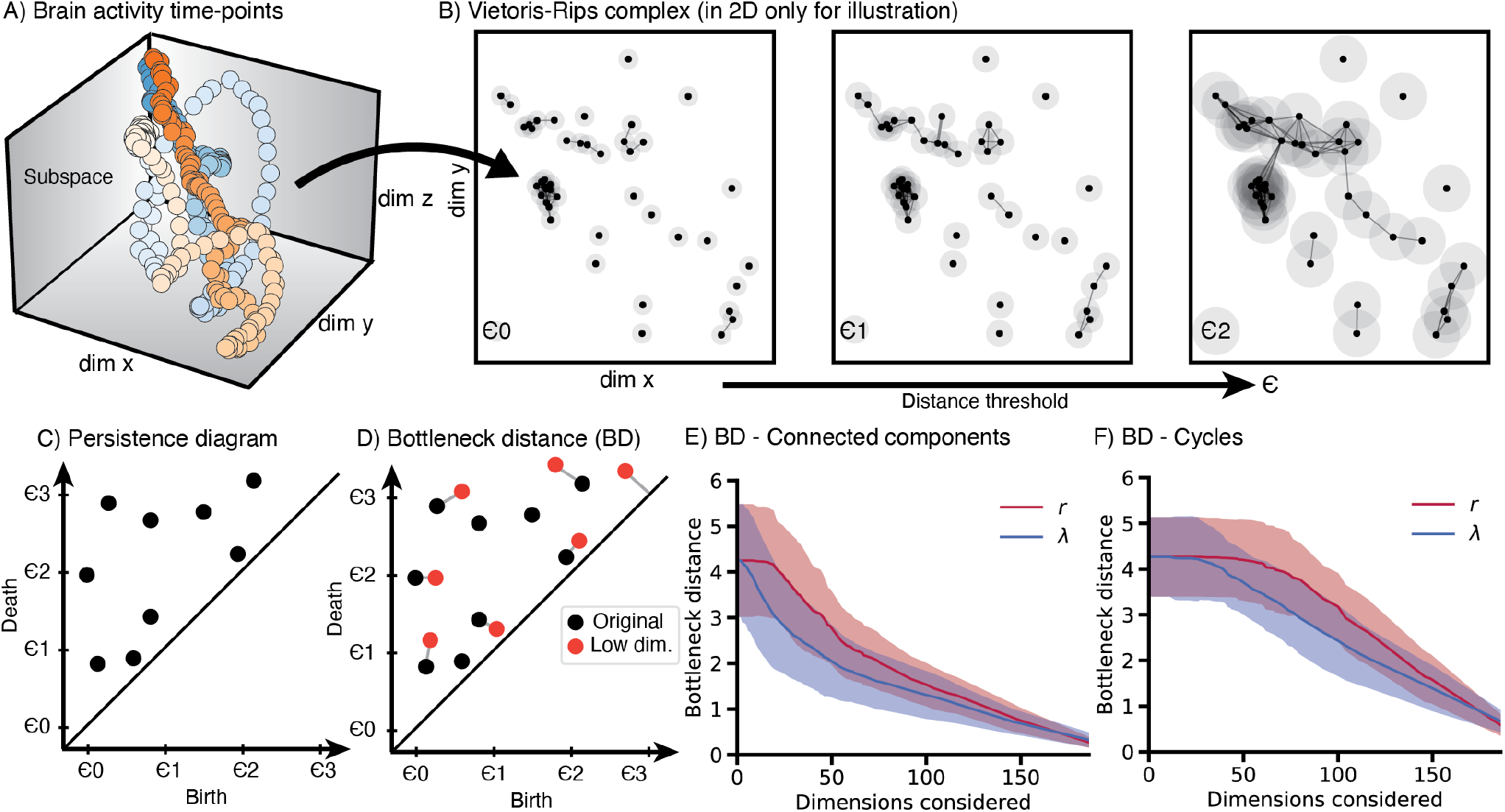
Topological features of the brain activity signal in different coordinate systems. **(A)** Two trajectories of brain activity (corresponding to two different stimuli) are represented in a low-dimensional subspace (here *d* = 3 for illustration purposes) **(B)** The Vietoris-Rips Complex (VRC) is constructed from the data points in **(A)** (in this example, only two dimensions are shown for illustration). The VRC is generated for each trajectory individually in the following manner: for a given distance threshold *ϵ*, connections or edges between data points smaller than the threshold are established (see *Methods: Persistence Diagrams and Bottleneck distance*). The number of simplicial complexes (connected components and cycles) that appear (birth) or disappear (death) when increasing *ϵ* is registered in the persistence diagram **(C)**. The persistence diagram summarizes the topology of the set of data points, i.e., the shape of the trajectory. **(D)** The distance between any two persistence diagrams can be computed with the bottleneck distance. **(E)-(F)** The bottleneck distance between the persistence diagrams of the original coordinate system (219 brain areas) and the reduced subspaces is shown (atlas-based and connectome spectrum). This distance is computed for connected components **(E)**, and for cycles **(F)**.

Following that rationale, we analyzed how the shape of the trajectory, in terms of topological features, was maintained when changing the number of dimensions in each coordinate system. We used the same criteria for reducing the dimensionality as before (dimensions were kept based on the energy they captured). The set of data points in the *d*-dimensional state-space (i.e., the signal, see Fig. 5A) was used to generate a Vietoris-Rips complex (*38*) (VRC, Fig. 5B). The VRC defines the connectivity among data points based on their proximity in the state-space spanned by *d* dimensions, for a varying distance threshold ϵ. For each distance threshold *ϵ*, the persistence diagram records the appearance (birth) or disappearance (death) of topological features (Fig. 5C). Once we summarized the shape of the brain activity trajectory in the two coordinate systems, we quantified how the persistence diagram changed when decreasing the dimensionality for each *d* in [1, *D*]. If a coordinate system conferred a good low-dimensional representation of the signal, we would expect to see that the persistence diagram of the subspace was very similar to the persistence diagram of the original high-dimensional representation. A bad low-dimensional representation would show a very different persistence diagram, as the important topological features would not be maintained. We used the bottleneck distance (*39*) (see *Methods: Persistence Diagrams and Bottleneck distance*) to quantify the distance between different persistence diagrams (see Fig. 5D). The results are shown in Fig. 5E-F. When the dimensionality of the subspace was small (i.e., *d* is less than 50 dimensions), both the atlas-based signal and its connectome spectrum failed to capture the topology of the original signal. However, as the dimensionality increased, the bottleneck distance in both coordinate systems decreased, indicating that the topological features were started to be accounted for. Importantly, the connectome spectrum representation of the signal better maintained, in comparison with the common brain atlas signal representation, the topology for the same number of dimensions (*p*<.001, Wilcoxon Rank-sum two-sided test). These results show that the fast temporal dynamics of brain activity are better preserved in a low-dimensional subspace defined by connectome harmonics.

## Discussion

A canonical signal representation is a representation that is the simplest or optimal in some aspect (*40*) (not to be confused with linear canonical transform (*41*)). In this work we have provided empirical evidence that the coordinate system spanned by the connectome harmonics has advantageous properties. When compared to a common area-based approach, connectome spectral analysis confers a sparser and more informative signal representation of large-scale brain activity dynamics, in the statistical, functional and topological sense.

### The connectome spectrum of brain activity is sparse

With the continuous advances in neurotechnology, spatial and temporal resolution of neural activity measurements are steadily improving. When analyzing noisy high-dimensional data, transformations into sparse signal representations have two main advantages: since the degrees of freedom are low, the interpretation and model generation is more tractable, and the robustness to noise is improved (*21*). These properties make compact coordinate systems optimal to construct efficient regularizers for regression and classification models, i.e., for brain decoding. We have shown that the signal is more compressible using a connectome spectral representation than using either the original signal representation by brain areas or surrogate connectome harmonics. This confers compelling evidence that the connectome spectrum decomposes brain activity in biologically relevant parts, adding one more piece of evidence that brain activity is shaped by macroscopic structural connectivity. Furthermore, the fact that connectome spectral dimensionality reduction performs equally or better than data driven approaches (PCA and ICA) suggests that the underlying description is relevant to macroscopic brain activity, as it captures the major axes of variability in the data. Hence, that the brain activity signal is an addition of smooth variations over structurally defined networks at small number of different scales (connectome harmonics) rather than a simple addition of the activity in individual brain regions. The plausibility of this hypothesis is strengthened by the positive linear relationship between the compactness of the connectome spectrum and the power of the signal (see Fig. 2C). For illustration, imagine a graph in which we inject a signal through one of the nodes. If the input signal has a strong power, the signal will diffuse along the whole graph following its edges, inducing strong coupling among distance nodes that will be captured by few low-frequency harmonics, and thus it will be compact. Conversely, if the injected signal is weak, the diffusion will not reach further than the first neighbors. This later activation pattern will take the shape close of a Dirac pulse, which translates into a flat spectrum, i.e., a non-compact connectome spectral representation.

In a previous article, we have shown that the process of face perception involves the greater activation of certain structural networks over single brain areas (*13*). Here we have observed different temporal regimes at which the connectome spectrum compactness is significantly higher than the compactness of the signal defined at the brain areas, and others at which they are equal. At the time of high-level face processing, at around 170 ms, the compactness of the signal is similar in both coordinate systems. This is due to the concentration of the activity into the brain regions around the FFA. Accordingly, the compactness of the signal in the original coordinate system increases, and the signal spectrum compactness decreases, i.e., the signal spectrum is more broadband. Despite this, when compressing the signal to 95% of the original, the reconstructed signal is globally better when compared to brain atlas-based compression. Fig. 2E shows how the signal spectrum compression preserves much more information than the compression in the original signal domain (see also Fig. S2D for the reconstruction at two other time-points). These results can be interpreted in the light of established mechanisms of face perception, which undergoes two pathways in the brain. The first pathway, which is the faster one (taking around 100 ms), is related to the processing of low-level spatial features. In this pathway, sub-cortical structures quickly activate and relay to other cortical regions, such as prefrontal areas (*42*). This fast and integrative process, would ideally be captured by few smooth harmonics, in a compact manner, as shown in the first significant peak in Fig. 2D. The second pathway, related to processing of higher-level features which help identify the face, activates the ventral pathway, and peaks at 170 ms in the FFA. This coarse activation pattern is reflected by the compactness deflection around 170 ms (*26*). After 170 ms, once the face image has been perceived, the participants engage a decision-making process during which the brain activity seems to be dominated by large scale structural networks again, i.e., low frequency harmonics. Then, between 300-600 ms after the stimulus presentation, the dynamics return to their baseline profile, dominated by higher-frequency harmonics. Such profile of compactness dynamics should be expected in any kind of visual experiment. We have replicated this result in a visual motion detection experiment (see Fig. S4). This was expected given that visual networks closely match the patterns defined by low-frequency harmonics (*11*), while a large number of harmonics of higher frequency are needed to compose networks associated to higher cognitive functions.

It is also important to take into consideration how the noise component of the signal is distributed in different coordinate systems. Before stimulus presentation, the recorded activity consists on the average brain activity across trials with very different brain states. Given that the time of the stimulus presentation is randomized, the brain signal prior to the stimulus that is left, i.e., the residual noise, is not associated to any particular dimension (brain area or connectome harmonic), being rather evenly distributed across dimensions. As indicated by a smaller compactness of the pre-stimulus signal in its connectome spectrum, noise co-varies less with connectome harmonics than with brain areas. This suggests that the brain activity signal representation in its connectome harmonics is more noise-independent than in the traditional atlas-based representation.

In summary, in this section we have shown that the brain activity signal is more compact when represented as an activation of connectome harmonics, or, in other words, that the brain activity signal is sparser in its connectome spectrum.

### Broadcasting dynamics are characterized by the connectome spectral content

Besides showing appealing statistical properties, the connectome spectrum seems to be a canonical basis for brain activity signal representation, as this later can be more easily explained by connectome harmonics than just by isolated brain areas. In fact, the connectome spectral power density follows a λ^−1^ decay, which indicates that, overall, the signal is smooth on the graph. A smooth signal on the graph is energy efficient, and informs us that the connections that are present in the structural connectivity are also present in the functional signal (*43*). In this paper, we have shown that the functional connectivity dynamics can be estimated without complex models simply by measuring at each time-point the contribution of scaled and orthogonal networks, namely connectome harmonics. We proposed a method to characterize the broadcasting profile of the brain activity signal at each time-point, characterizing brain activity in terms of integration and segregation mechanisms with respect to the underlying structural network (*44, 45*). We have been inspired by the fact that functional integration and segregation are strongly related to connectome harmonics (*20*), with low-order harmonics being associated to integration and higher-order harmonics being associated to segregation mechanisms. With this approach applied to visual evoked potentials in a face perception experiment (and in a motion perception experiment in the supplementary material), we have discovered that integration and segregation follow consistent fast dynamics across all participants, and are closely related to the compactness of the signal in connectome spectrum (see Fig. 2D). During high-order visual processes, such as the integration of visual features into a whole object, or the decision making that follows, the brain activity signal is dominated by low-frequency harmonics. During that period, the brain is in an integrated configuration, and there is an increase in signal compactness. Conversely, during segregated processing, i.e., the initial parallel processing of low-level features, or the peak activation of FFA during face perception, high-order harmonic components add in, collectively reducing the signal compactness.

### The structure of the brain shapes its dynamics

So far, we have shown that connectome spectral decomposition not only provides us with a robust and sparse representation for brain activity, but also enables a natural description of the broadcasting status of the signal, indicating whether the brain in integration or segregation mode. These can be regarded as some general properties of the signals defined on the graph. We wanted to evaluate whether connectome spectral analysis can show any added value to the study of brain activity dynamics. In the context of theoretical neuroscience, it can provide a powerful methodology to represent the dynamics of brain activity as a trajectory in its state-space (*46*) (see Fig. 4A and 5A). In this context, the working hypothesis is that high-level neural computation involves the formation of low-dimensional structures in the state-space known as attractors (*47, 48*). Here, we have studied low-dimensional structures spanned by areas and connectome harmonics. Our results show that functionally relevant properties of the signal are characterized with more sensitivity when the state-space is defined by the signal’ s connectome spectrum, rather than by the activity of areas. More specifically, we have characterized two different properties of the signal. On the one hand, we have shown that the distance between the trajectories corresponding to brain activity processing of two different types of information (low-level vs. high-level visual stimuli features) is higher in the subspace of the connectome spectrum. On the other hand, we have investigated another important property of the state-space: its contribution to the shape of the signal intrinsic structure (*35*). Using topological data analysis (*36, 37*), we have quantified the contribution of each state-space to the overall shape of the brain activity trajectory. The results of this last analysis indicated that the connectome spectrum representation of the signal captures the most important features of the dynamics with a smaller number of dimensions.

### Limitations and future directions

There are some important limitations in our study. The most important of them is associated to the use of EEG data. The spatial smoothness of the inverse solution used could bias the estimated coupling between structure and function. This limitation is made obvious by the fact that there is one type of connectome spectrum configuration that we have not observed in our data: the activation of single, or very few high-frequency harmonics during the visual evoked activity (see Fig. 1F). In our datasets, these components activate very weakly. The relatively smooth nature of the reconstructed electrical activity imposes that very high-frequency connectome harmonics will always have small weight in the signal. However, the smoothness of the source reconstruction does not completely account for the prominence of the low-frequency components, which are spatially much smoother (see Fig. S1).

A related limitation is the general difficulty of the Fourier transform to pick up transients in the graph, such as a Dirac delta functions, in a compact manner. This is the same in the case of spatial and temporal Fourier analysis. Such transient patterns are better captured by the signal representation based on brain areas (which are Dirac delta functions). A possible solution to be investigated in future work could be the use graph wavelets (*49*) instead of connectome harmonics, as they excel at picking up transient signals in a compact manner.

Our results suggest that the shape of the structural connectivity plays a very important role in the signal decomposition. Interesting future directions could be to study how the inter-individual variability on the structural connectivity modulates the spectral representation of the signal. This approach would very challenging, given that different graph-structures would provide different connectome harmonics, and the comparison between subjects would not be straight forward. Similarly, future directions could point towards hallmarks of brain disorders in which the structural connectivity is affected.

A potential different line of research to follow-up on this work, would be the study of the time-vertex joint power-spectral density (see (*25*)). In that direction, the joint signal representation of spatial and temporal modes could be related to the theories of communication through coherence (*50*) and brain oscillations (*51*). In this sense, it would be very interesting to investigate the role of the connectivity measures used to define the edges in the structural connectivity matrix. Here we used the number of fibers connecting every pair of brain areas (estimated from tractography). The connectome harmonics would have been different if we used the average fiber length instead. This last approach could be useful to investigate the role of delays in the functional connectivity.

Finally, further validation is needed in different contexts, to verify whether the conclusions and hypotheses presented hold for a larger variety of tasks.

## Materials and Methods

### Data acquisition and pre-processing

High-density EEG was recorded at 2048 Hz using a 128-channel Biosemi Active Two EEG system (Biosemi, Amsterdam, The Netherlands;) at the Fribourg Cantonal Hospital, Fribourg, Switzerland. A total of 20 healthy participants (17 females, mean age: 23, age range: 20-29 years) were recorded while performing a visual discrimination task. A good signal quality was guaranteed by keeping the offset between the active electrodes and the Common Mode Sense -Driven Right Leg (CMS-DRL) feedback loop under a standard value of ±20 mV. After each participant session, individual 3D electrode positions were digitized with an ultrasound motion capture system (Zebris Medical GmbH). This open dataset (*52*) has been used in previous publications (*13, 24*). The visual stimuli consisted on images of faces or scrambled versions of the same images (*53*). After a 200 ms image presentation, participants responded, by pressing one of two buttons on a response box, whether they had seen a face or a scrambled image. One participant was excluded due to motion artifacts, leaving 19 datasets for analysis. Data were pre-processed using EEGLAB v14.1.1 (*54*) (*sccn*.*ucsd*.*edu/eeglab/index*.*php*). Down-sampling to 250 Hz (anti-aliasing filter: cut-off frequency of 112.5 Hz; transition bandwidth of 50 Hz) and local detrending (high-pass filter at 1 Hz, EEGLAB PREP plugin) were applied (*55*). Trials were extracted from 1500 ms before the stimulus presentation until 1000 ms after. Line and monitor noise (at 50 and 75 Hz, respectively, as well as harmonics of these frequencies) were removed by spectral interpolation (*56*). Bad trials (22 ± 36 out of 600 per subject) were removed and bad channels (15 ± 9 out of 128 per subject) marked via visual inspection. The remaining physiological artifacts (eye blinks, horizontal and vertical eye movements, muscle potentials) were removed using FastICA by first marking bad ICs using Multiple Artifact Rejection Algorithm (MARA) as implemented in EEGLAB (*54*). The previously identified bad channels were not included in this step. Finally, bad channels were interpolated using the nearest neighbor spline method as implemented in EEGLAB, and data were re-referenced to the common average before being globally z-scored.

### Structural MRI

T1-weighted images were obtained from the same subjects as magnetization prepared rapid-gradient echo (MPRAGE) volumes using a General Electrics Discovery MR750 3T MRI scanner and a COR FSPGR BRAVO pulse sequence with flip angle = 9°; echo time = 2.81 ms, repetition time = 7.27 ms, inversion time = 0.9 s, slice thickness = 1 mm, head first supine. Connectome mapper 3 (*57*) v3.0.0-RC1 with Freesurfer 6.0.1 were used to perform the segmentation of the MPRAGE volume into gray and white matter, as well as the parcellation of gray matter brain regions of interest according to the Lausanne 2008 multiscale parcellation (*9*). Connectome Mapper 3 and Freesurfer are available at *github*.*com/connectomicslab/connectomemapper3* and *surfer*.*nmr*.*mgh*.*harvard*.*edu/fswiki/FreeSurferWiki*.

### Diffusion MRI

To create the structural connectivity matrix, we obtained a consensus connectome from an online available dataset consisting on 70 healthy participants (*58*) available at *zenodo*.*org/record/2872624*; mean age 29.7 years, range 18.5-59.2 years; 34 females), scanned in a 3-Tesla MRI scanner (Trio, Siemens Medical, Germany) with a 32-channel head-coil. A diffusion spectrum imaging (DSI) sequence (128 diffusion-weighted volumes and a single b0 volume, maximum b-value 8,000 s/mm2, 2.2×2.2×3.0 mm voxel size) was applied, and DSI data were reconstructed following the protocol described in (*59*). A magnetization-prepared rapid acquisition gradient echo (MPRAGE) sequence sensitive to white/gray matter contrast (1 mm in-plane resolution, 1.2 mm slice thickness) was also applied, and gray and white matter were segmented from the MPRAGE volume using Freesurfer and Connectome Mapper 3 (*57*). Then, individual structural connectivity matrices were estimated by deterministic streamline tractography on the reconstructed DSI data, by seeding 32 streamlines per diffusion direction in each white matter voxel (*59*). The number of fibers found between each voxel at the gray matter/white matter-interface was summed within each brain area given by the same parcellation used for the structural data.

A consensus group-representative structural brain connectivity matrix was generated from the connectomes of 70 healthy participants connectomes using the method introduced in (*60*). This method selects a recurrence threshold that preserves the connection density of each single subject connectome. The connection density is preserved independently for intra- and interhemispheric connections, allowing a more inter-hemispheric connections to be kept in the group estimate, in comparison to simple connectome average across subjects. The resulting connection density in the group connectome is set to 25%.

### Source reconstruction and atlas-based time-courses

Grey matter (source) signal reconstruction of the scalp EEG signals was performed using individual realistic head conductor models, based on the tissue segmentation of the individual structural images and the EEG electrode positions. Both forward solutions, using the Locally Spherical Model with Anatomical Constraints (LSMAC) and inverse solutions, using the Local Autoregressive Average (LAURA) method, were implemented with the CARTOOL toolbox (*61*) (freely available at *https://sites.google.com/site/cartoolcommunity/*).

Using the inverse solution matrix, we projected the EEG data of each participant into approximately 5000 dipole locations uniformly distributed in the gray matter. Apart from a magnitude, each dipole has a three-dimensional orientation. The time-series for each brain area was extracted from dipoles whose location overlapped with that region. Using the singular value decomposition method (*24*), we estimated the orientation of maximal variance during the time window defined from 120 to 500 ms after stimulus onset. Each dipole in the brain area was then projected to the estimated orientation, and the average magnitude of the projection across dipoles was taken as the brain area value for each time-point.

### Graph Signal Processing

#### Connectome harmonics and Graph Fourier Transform

The connectome harmonics are the eigenfunctions of the structural connectivity graph Laplacian. Given a structural connectivity graph *𝒢*(*D, E*) of *D* nodes and *E* edges, we define the graph Laplacian as:

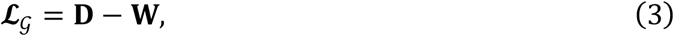

where **D** is the degree matrix and **W** is the weight matrix. Here we used the normalized graph Laplacian, which is defined as:

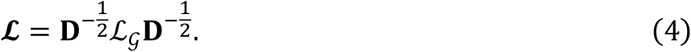

The connectome harmonics are obtained by the eigendecomposition of the normalized graph Laplacian:

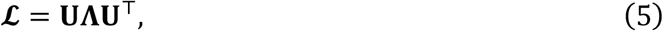

where [Λ] _*dd*_ = λ_*d*_ for *n* = [1, …, *D*] are the eigenvalues of the graph Laplacian ordered according to smoothness, and are associated to the *d*-th eigenvector (connectome harmonic) contained in the *d*-th column of *U*. We formulate the graph Fourier transform 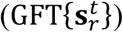 of the brain activity defined in the atlas-based coordinate system at a given moment in time 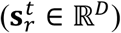 as following:

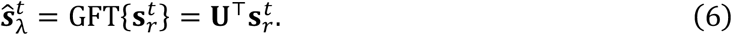

The GFT linearly maps the original signal into the connectome harmonics. To project back the signal to the atlas-based coordinate system, we define the inverse graph Fourier transform iGFT as:

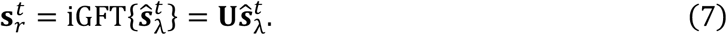

These operations were implemented using the Graph Signal Processing toolbox in Python available at *https://pygsp.readthedocs.io/en/stable/*.

#### Graph power spectral density

The graph power spectral density describes the amount of energy present in each connectome harmonic during a graph time-varying signal. We compute the normalized graph power spectral density for each participant by: first transforming the evoked signal (i.e., the average signal across trials) into its connectome spectrum through the GFT. Then normalizing the connectome spectrum of the evoked signal by the standard deviation in time. Finally, the power is defined as the mean squared signal spectrum across time.

#### Graph spectrum dichotomization

The graph spectrum dichotomization was first introduced in (*34*). First, we decompose the brain activity signal in low-frequency and high-frequency connectome harmonics parts. These two signals are computed by multiplying the graph Fourier transformed coefficients by the indicator function to the first T lowest eigenvalues and to the *D* − T highest eigenvalues, where λ_*T*_ is the threshold graph-frequency that divides the graph energy spectrum in two halves:

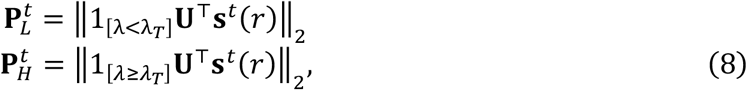

These power time-courses inform about the amount of energy of the original signal contained in the lowest- and highest-end of the graph spectrum for each time-point.

#### Surrogate harmonics

A thousand randomized graphs were obtained using the Brain Connectivity Toolbox (*62*) function *null_model_und_sign*. The harmonics from these graphs were obtained using the above-mentioned approach, and are referred as surrogate harmonics in this manuscript.

### Analysis of compactness

#### Signal compactness in a given coordinate system

Signal compactness was measured using two metrics: The Person correlation between the original and the compressed signal, and the compression reconstruction error. These two metrics are based on first compressing the signal using:

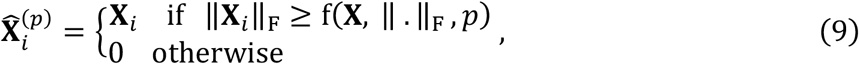

for **X** ∈ ℝ^*D,T,N*^ and **X**_*i*_ ∈ ℝ^*T,N*^ corresponding to the *T* samples and *N* trials of the *i*-th dimension out of *D* total dimensions (brain areas or harmonics), where ‖ · ‖_F_ is the Frobenius norm. f() is a function that returns the value of the *p*-th percentile in the distribution of the norms of all dimensions. This compression removes the dimensions with less amount of signal power throughout time and trials.

Given a signal **X**^(S)^ ∈ ℝ^*D,T,N*^, *D* being the dimensionality, *T* the number of samples, and *N* the number of trials, and 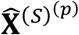 being the compressed signal, the reconstruction normalized mean squared error for a given percentile is computed as:

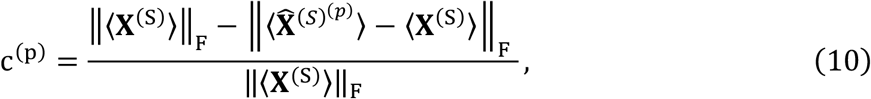

where ⟨ · ⟩ denotes the average across trials. Compression correlation is defined as:

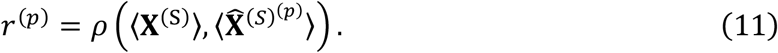

When computing compactness dynamics, we first define compression as:

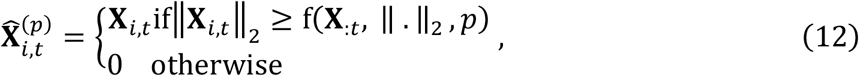

for **X**_*i,t*_ ∈ ℝ^*N*^ corresponding to the *N* trials of the *i*-th dimension at sample *t* of **X**, where ‖ · ‖_2_ is the ℓ_2_-norm. f() is a function that returns the value of the *p*-th percentile in the distribution of the norms of all dimensions. We then define compactness dynamics (*c*_*t*_) as the average reconstruction performance (1 -reconstruction error) of the compressed signal across all percentiles at each time-point:

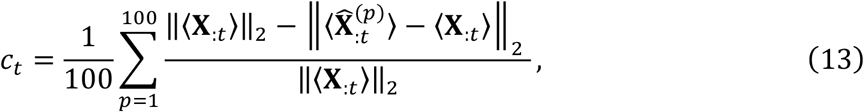

where ⟨ **X**_:*t*_⟩ ∈ ℝ^*D*^ (and 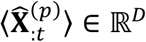) is the average pattern (across trials) of the original signal (compressed signal at a given percentile) across all trials at time *t*. Please note the differences to when measuring compactness in Eq. 10.

#### Compression effect on visual streams

We assessed whether the effect of compressing the signal was harmful at specific brain areas known to be involved on visual processing. For that reason, we defined the dorsal and ventral streams, and a group of early visual areas, by clustering some of the 219 regions of interest in scale 3 of the multi-scale Lausanne atlas parcellation (*23*). The choice of brain areas used is shown in Fig. S3, and was based on a previous study finding clusters corresponding to the different visual pathways (see Fig. S4 in the ref. (*9*)). Compactness metrics were computed only taking into account the signal on those areas. To do so, in the case of the connectome spectrum, the signal of brain region activity was uncompressed using a set of coefficients estimated by compression, by means of the iGFT defined in Eq.7.

#### Conditional probabilities

The conditional probability of the compactness of the evoked signal ⟨ **X**^(S)^⟩ ∈ ℝ^*D,T*^ in a given coordinate system S, given the signal power was computed as the following. First, we computed the compactness dynamics, using Eq. 12. We then removed the outliers (3 standard deviations) and computed the joint probability of signal compactness and power:

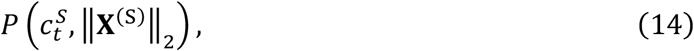

using the Gaussian kernel density estimation from the python SciPy toolbox (*63*). From the joint probability, we obtained the marginal probability of the signal power. The conditional probability was finally obtained as:

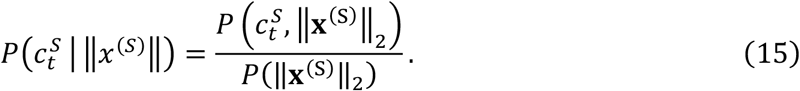

### Dimensionality analysis

#### Co-linearity of dimensions

Co-linearity of dimensions was measured by first computing the correlation matrices:

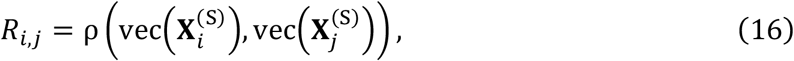

where 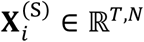 corresponds to the data in the *i*-th dimension of the coordinate system S, with *T* samples and *N* trials, and after vectorization 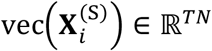.

#### Distance difference between different trajectories

We computed the distance between two trajectories for each time-point as:

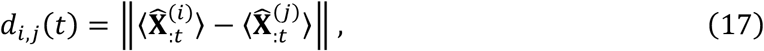

where 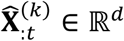 is the average pattern across the trials corresponding to the *k*-th condition at time *t*.

#### Low-dimensional embedding

We followed the dimensionality reduction approach based on the same criterion as the compression in Eq. 9. Namely, to reduce the dimension to a given percentile *p* we keep the *i*-th dimension if:

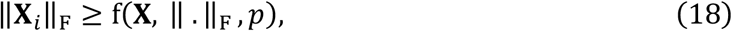

for **X**_*i,t*_ ∈ ℝ^*N*^ corresponding to the *N* trials of the *i*-th dimension at sample *t* of **X**, where ‖ · ‖_F_ is the Frobenius norm. f() is a function that returns the value of the *p*-th percentile in the distribution of the norms of all dimensions.

#### Persistence Diagrams and Bottleneck distance

To compute the persistence diagrams of the trajectories, we used the package Ghudi available at *http://gudhi.gforge.inria.fr*. Persistence diagrams were generated for connected components and cycles, for coordinate systems with varying number of dimensions, using the low-dimensional embedding explained above. Then we computed the bottleneck distance between the persistence diagrams of the original signal (219 areas) and the persistence diagram of the signal in the subspace of the different dimensions.

## Author contributions

Conceptualization: JRQ, KG, GP, PH

Data collection and pre-processing: DP, ST, MC, JC Investigation: JRQ, KG, DP, GP, PH

Visualization: JRQ Supervision: GP, PH, SV

Writing—original draft: JRQ, GP, PH

Writing—review & editing: JRQ, KG, DP, ST, MC, SV, GP, PH

## Data and materials availability

Links to all data and code are available in the main text.

