## Supplementary Figures for "The connectome spectrum as a canonical basis for a sparse representation of fast brain activity"

**This PDF file includes:**

Figs. S1 to S4

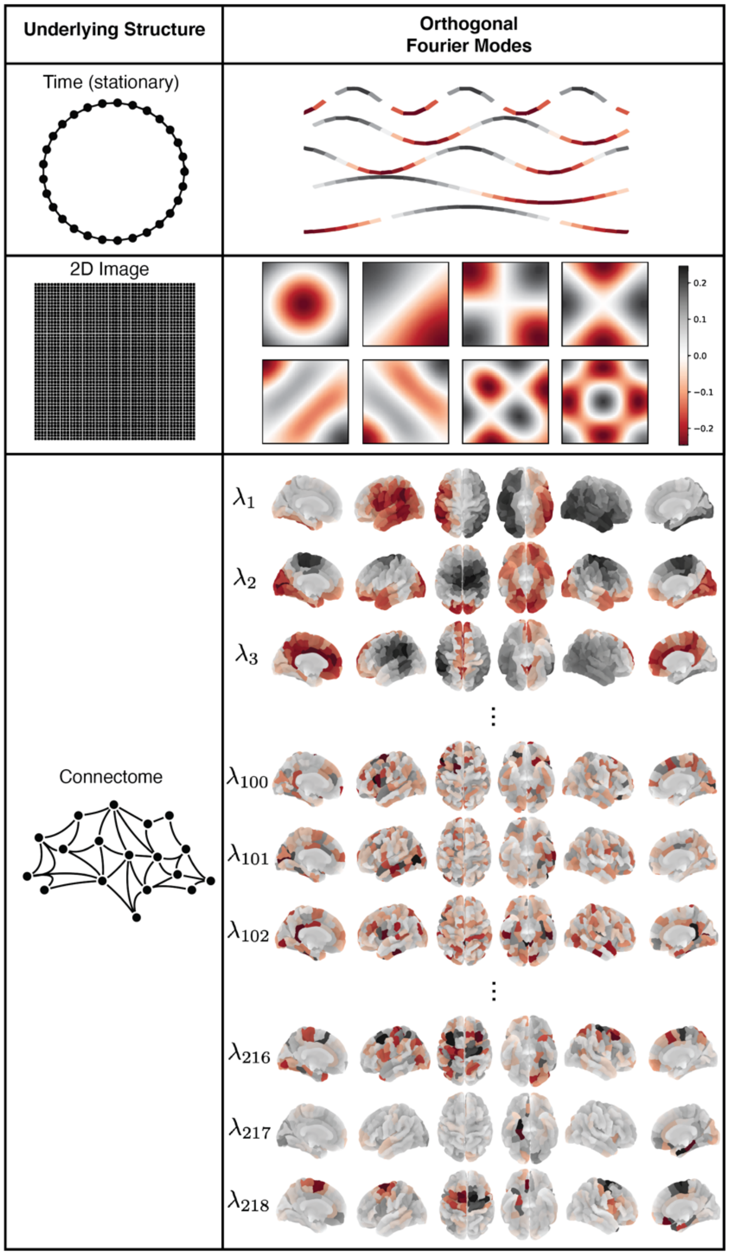


**Fig. S1.**
**The graph Fourier transform (GFT).** The GFT is a generalization of the Fourier transform to graph domain structures. When the underlying graph is a unitary circle, the GFT corresponds to the discrete FT (DFT), and the circular graph Laplacian eigenvectors correspond to sinusoids with different frequency. When the underlying graph is a 2D grid, the GFT corresponds to the discrete two-dimensional FT, widely used for image processing, and the its graph Laplacian eigenvectors correspond two-dimensional shown in the second row. When the graph is defined by the structural connectivity, the eigenvectors define the connectome harmonics. Here we show three harmonics associated with a small graph Laplacian eigenvalue, three harmonics from the middle of the spectrum, and three harmonics from the highest end of the spectrum.

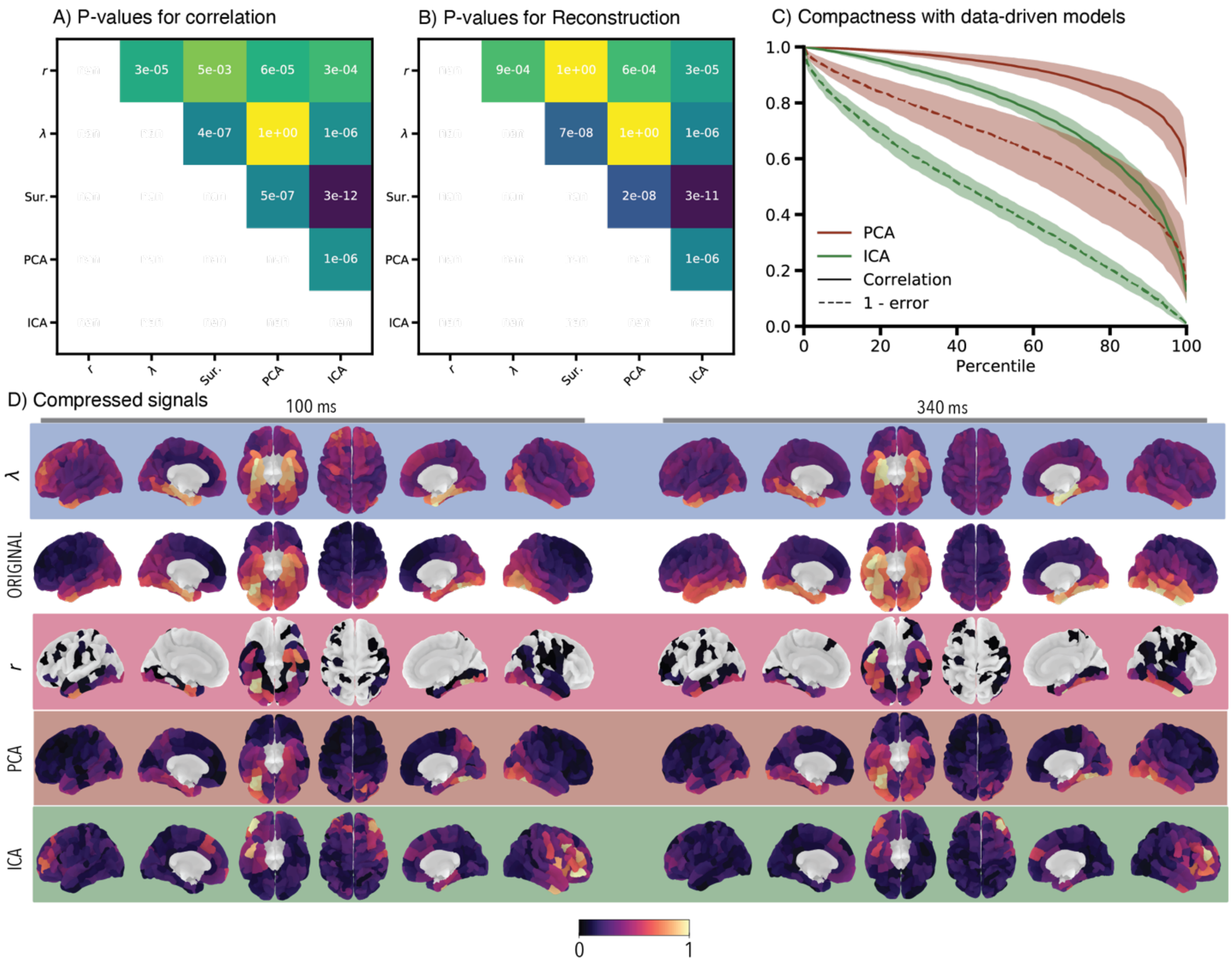


**Fig. S2.**
**Signal compactness in different coordinate systems.** This figure contains additional information for the figure 2 of the main manuscript. **(A)** Contains the p-values for the pair-wise Wilcoxon Ranksum statistical test of the median correlations, corrected for multiple comparisons with the Bonferroni method. **(B)** Same as **(A)** but for the reconstruction error. **(C)** Compactness profile for PCA and ICA representation of the signal (similar to plot in Fig 2A). **(D)** Reconstruction maps at two different time-points after the signal is compressed in the different coordinate systems (at a level of compression of 5%).

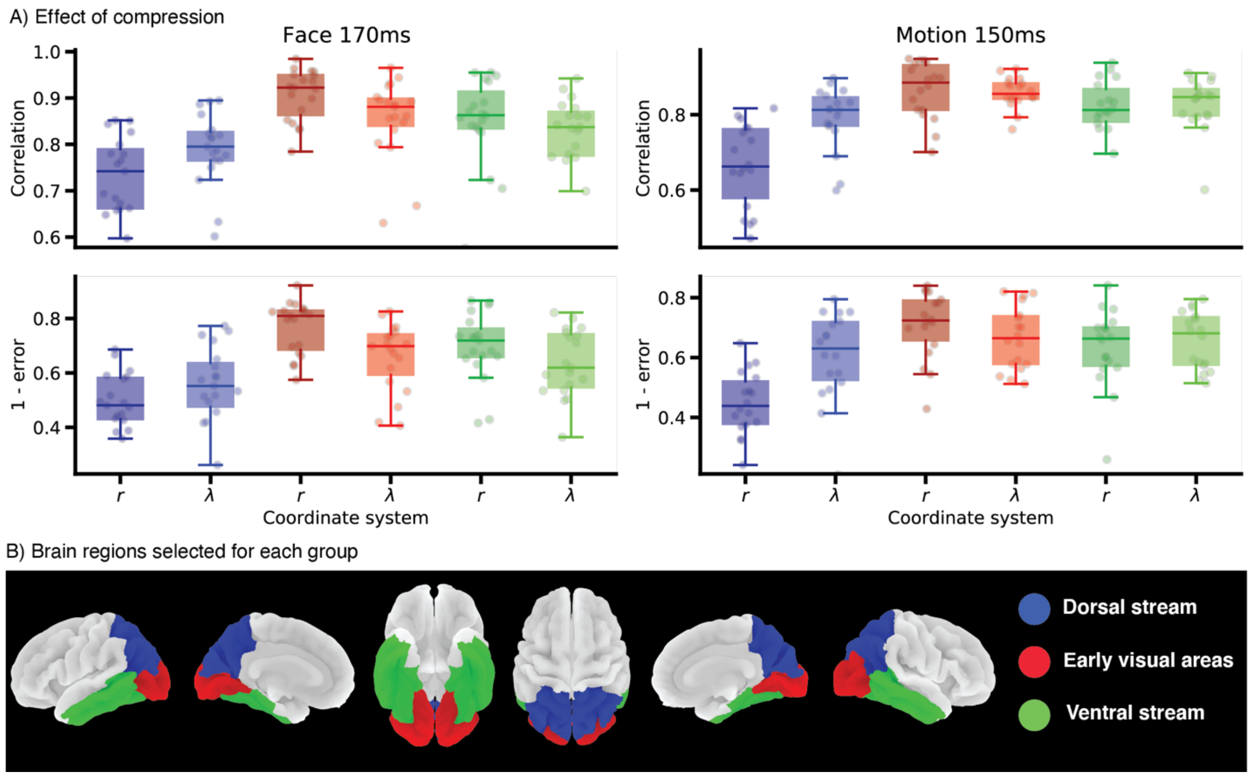


**Fig. S3.**

**Signal compactness in visual systems for known perceptual processes.** For the three visual systems shown in **(B)**, **(A)** shows the compactness performance (similar to Fig.2B) for two well-known perceptual processes (face perception at 170 ms after stimulus presentation, and motion perception at 150 ms after stimulus presentation). r refers to the atlas-based signal representation, and λ refers to the connectome spectrum signal representation.

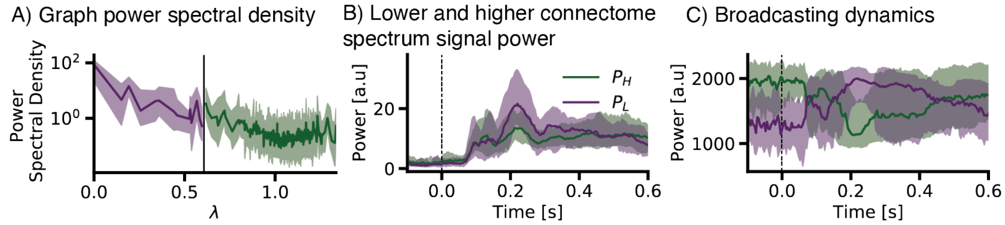


**Fig. S4.**
**Validation of broadcasting dynamics in a different dataset.** This figure shows the same plots as Fig. 3 in the main manuscript but for a motion processing visual experiment.
